# Mechanism of negative regulation of NF-κB by N4BP1

**DOI:** 10.1101/2020.10.21.349357

**Authors:** Hexin Shi, Lei Sun, Ying Wang, Aijie Liu, Xiaoming Zhan, Xiaohong Li, Miao Tang, Priscilla Anderton, Sara Hildebrand, Jiexia Quan, Sara Ludwig, Eva Marie Y. Moresco, Bruce Beutler

## Abstract

Many immune responses depend upon activation of NF-κB, a key transcription factor in the elicitation of a cytokine response. Here we demonstrate that N4BP1 inhibited TLR-dependent activation of NF-κB by interacting with the NF-κB signaling essential modulator (NEMO, also known as IκB kinase γ) to attenuate NEMO-NEMO dimerization or oligomerization. The UBA-like (ubiquitin associated-like) and CUE-like (ubiquitin conjugation to ER degradation) domains in N4BP1 mediated the interaction with the NEMO COZI domain. Both *in vitro* and in mice, *N4bp1* deficiency specifically enhanced TRIF-independent (TLR2, TLR7, or TLR9-mediated), but not TRIF-dependent (TLR3 or TLR4-mediated), NF-κB activation leading to increased production of proinflammatory cytokines. In response to TLR4 or TLR3 activation, TRIF caused activation of caspase 8, which cleaved N4BP1 distal to residues D424 and D490 and abolished its inhibitory effect. *N4bp1*^*-/-*^ mice also exhibited diminished numbers of T cells in the peripheral blood. Our work identifies N4BP1 as an inhibitory checkpoint protein that must be overcome to activate NF-κB, and a TRIF-initiated caspase 8-dependent mechanism by which this is accomplished.

---

The inducible transcription factor NF-κB plays a key role in the development and function of the immune system by activating or repressing hundreds of genes that carry out the cellular response to specific stimuli^1^. Canonical activation of NF-κB requires its release from the inhibitory IκB proteins that normally retain NF-κB in the cytoplasm^2^. This occurs when the IκB kinase (IKK) complex phosphorylates IκB proteins resulting in their subsequent K48-ubiquitination and degradation by the proteasome. In this role, the IKK complex serves as the gatekeeper for NF-κB activation and as such is subject to numerous modifications that modulate its activity. Composed of three subunits, the kinases IKKα and IKKβ and the regulatory subunit NEMO, the IKK complex is activated by combinations of phosphorylation (on IKK), ubiquitination (on NEMO), and sumoylation (on NEMO) events, and negative regulators target these modifications to either remove or prevent them^3, 4^. Persistent NF-κB activation may lead to pathologic consequences such as chronic inflammation or autoimmunity^5^.

Although receptor-proximal signaling pathways differ, inflammatory stimuli such as tumor necrosis factor (TNF), interleukin (IL)-1, and ligands for the Toll-like receptors (TLRs) all activate NF-κB, impinging on the IKK complex to do so. NF-κB then enters the nucleus to activate the transcription of many proinflammatory genes including those encoding TNF and IL-6. We carried out a forward genetic screen to discover proteins that mediate TLR signaling, testing peritoneal macrophages from *N*-ethyl-*N*-nitrosourea (ENU)-mutagenized mice for altered TNF responses to stimulation with TLR ligands. TLR activation results in the recruitment of up to four adaptor proteins, MyD88 (myeloid differentiation 88), MAL (MyD88-adaptor like), TRIF (Toll-interleukin 1 receptor (TIR) domain-containing adaptor inducing IFN*β*) and TRAM (TRIF related adaptor molecule), which assemble distinct signaling complexes depending on the combination of adaptors utilized. All TLRs except TLR3 depend upon MyD88 for full signaling activity whereas TRIF is utilized only by TLR3 and TLR4^1, 6, 7^.

We identified several mutations in the neural precursor cell expressed, developmentally down-regulated 4 binding protein 1 gene (*N4bp1*) that caused elevated TNF production by macrophages stimulated with ligands for TLRs other than TLR3 and TLR4. Here, we describe mechanistic studies showing how N4BP1, initially identified as a NEDD4 binding protein^8^, inhibits NF-κB activation downstream from all TLRs except TLR3 and TLR4. We also demonstrate how TLRs that recruit TRIF overcome the inhibitory effect of N4BP1 on NF-κB.

## Results

### Multiple *N4bp1* mutations enhance TRIF-independent TLR signaling

Peritoneal macrophages from third-generation (G3) descendants of ENU-mutagenized mice were assayed for TNF secretion after stimulation with TLR ligands. Three distinct mutations of *N4bp1* (*stash, winter*, and *acorn*) (Fig. 1a) in three ancestrally unrelated pedigrees were associated with increased production of TNF in response to R848, a TLR7 ligand (Fig. 1b,c and Supplementary Fig. 1a-c). The *acorn* (*ac*) allele contained the nonsense mutation Q135* and was a predicted null allele. We detected no N4BP1 protein in *N4bp1*^*ac/ac*^ macrophages, consistent with nonsense-mediated decay of the transcript (Fig. 1d).

**Figure 1.**
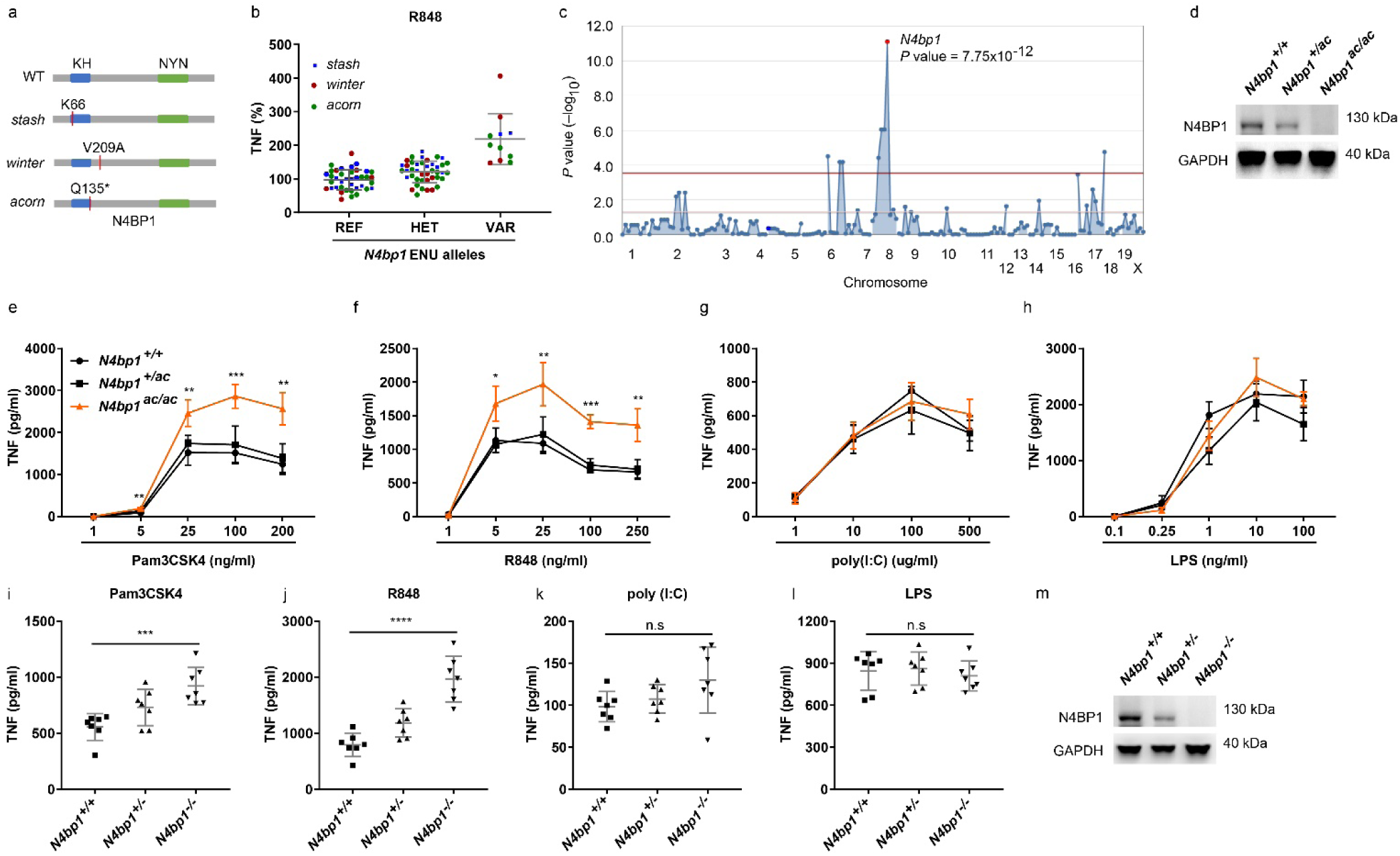
*N4bp1* mutations enhanced TRIF-independent TLR signaling. **a**, N4BP1 domain structure and positions of ENU-induced mutations (red lines). **b**, Relative concentration of TNF in the culture medium of peritoneal macrophages harvested from mice of the *stash* (blue), *winter* (red), and *acorn* (green) pedigrees 4 h after stimulation with 20 ng/ml of R848. REF, *N4bp1*^*+/+*^; HET, *N4bp1*^*+/mutation*^; VAR, *N4bp1*^*mutation/mutation*^, where “mutation” represents the *walnut, winter*, or *acorn* allele. **c**, Manhattan plot showing *p* values of association between the phenotype of elevated TNF production in response to R848 and mutations identified in the three pedigrees in (b) calculated using a recessive model of inheritance. The −log10 *p* values were plotted versus the chromosomal positions of mutations. Horizontal lines indicate thresholds of *p* = 0.05 with (red) or without (purple) the Bonferroni correction. The *p* value for linkage of *N4bp1* mutations with the elevated TNF production is indicated. **d**, Immunoblot analysis of N4BP1 and GAPDH in *N4bp1*^*+/+*^, *N4bp1*^*+/ac*^ and *N4bp1*^*ac/ac*^ peritoneal macrophages. **e-h**, TNF concentration in the culture medium of *N4bp1*^*+/+*^, *N4bp1*^*+/ac*^ and *N4bp1*^*ac/ac*^ peritoneal macrophages treated with Pam3CSK4 (**e**), R848 (**f**), poly(I:C) (**g**), and LPS (**h**). n=4 mice per genotype. **i-l**, TNF concentration in the culture medium of *N4bp1*^*+/+*^, *N4bp1*^*+/-*^ and *N4bp1*^*-/-*^ peritoneal macrophages treated with 40 ng/ml Pam3CSK4 (**i**), 20 ng/ml R848 (**j**), poly(I:C) (**k**), and LPS (**l**). **m**, Immunoblot analysis of N4BP1 and GAPDH in *N4bp1*^*+/+*^, *N4bp1*^*+/-*^ and *N4bp1*^*-/-*^ peritoneal macrophages. Each symbol (**b**,**i-l**) represents an individual mouse. * *P* < 0.05, \*\**P* < 0.01, \*\*\**P* < 0.001, \*\*\*\**P* < 0.0001 (One-way ANOVA). n.s, not significant. Data are representative of two (**e-h**) or three (**i-l**) independent experiments (mean ±s.d. in **b**,**e-l**).

Further analysis of responses to TLR stimulation demonstrated dose-dependent increased production of TNF by *N4bp1*^*ac/ac*^ macrophages stimulated with Pam3CSK4 (TLR2-TLR1 ligand, Fig. 1e) or R848 (Fig. 1f). However, *N4bp1*^*ac/ac*^ macrophages produced similar amounts of TNF as *N4bp1*^*+/+*^ macrophages when stimulated with poly(I:C) (TLR3 ligand, Fig. 1g) or lipopolysaccharide (LPS; TLR4 ligand, Fig. 1h). Peritoneal macrophages (Fig. 1i, j), bone marrow-derived macrophages (BMMs) (Supplementary Fig. 1e), and bone marrow-derived dendritic cells (BMDCs) (Supplementary Fig. 1f) from mice homozygous for a CRISPR-Cas9-targeted null allele of *N4bp1* (Fig. 1m) recapitulated the *acorn* phenotype, responding to Pam3CSK4 and R848 by producing excessive amounts of TNF and interleukin (IL)-6. Time course analysis of the cytokine response in *N4bp1*^*-/-*^ peritoneal macrophages supported elevated production rather than a merely accelerated response (Supplementary Fig. 1h). In contrast, the TNF and IL-6 responses to poly (I:C) and LPS were normal (Fig. 1k, l and Supplementary Fig. 1e, f). IFNα production induced by dsDNA stimulation was comparable between wild-type and *N4bp1*^*-/-*^ BMMs and BMDCs (Supplementary Fig. 1g). These findings indicate that TLR2-TLR1 and TLR7 signaling, but not TLR3 and TLR4 signaling, were enhanced in *N4bp1*^*-/-*^ macrophages and dendritic cells. Since TLR3 and TLR4 depend on the TRIF adapter protein to propagate signaling, whereas all other TLRs signal independently of TRIF, we concluded that N4BP1 inhibits TRIF-independent signaling.

Among tissues examined, the 893-amino acid N4BP1 protein was highly expressed in lymphoid organs, brain and ovary (Supplementary Fig. 2a). Cellular fractionation of peritoneal macrophages stimulated with R848 for up to 4 hours showed that N4BP1 was predominantly localized in the cytosol before and throughout the period of stimulation (Supplementary Fig. 2b). We also detected reduced numbers of CD4^+^ and CD8^+^ T cells in the peripheral blood of *N4bp1*^*-/-*^ mice (Supplementary Fig. 2c). The numbers of peripheral blood monocytes, neutrophils, and B cells were normal (Supplementary Fig. 2c). *N4bp1*^*-/-*^ mice appeared grossly normal and showed no internal anatomical abnormalities. The mice were not overtly suffering from autoimmunity since the concentration of anti-dsDNA antibodies in the serum was comparable with that in wild-type mice (Supplementary Fig. 2d).

**Figure 2.**
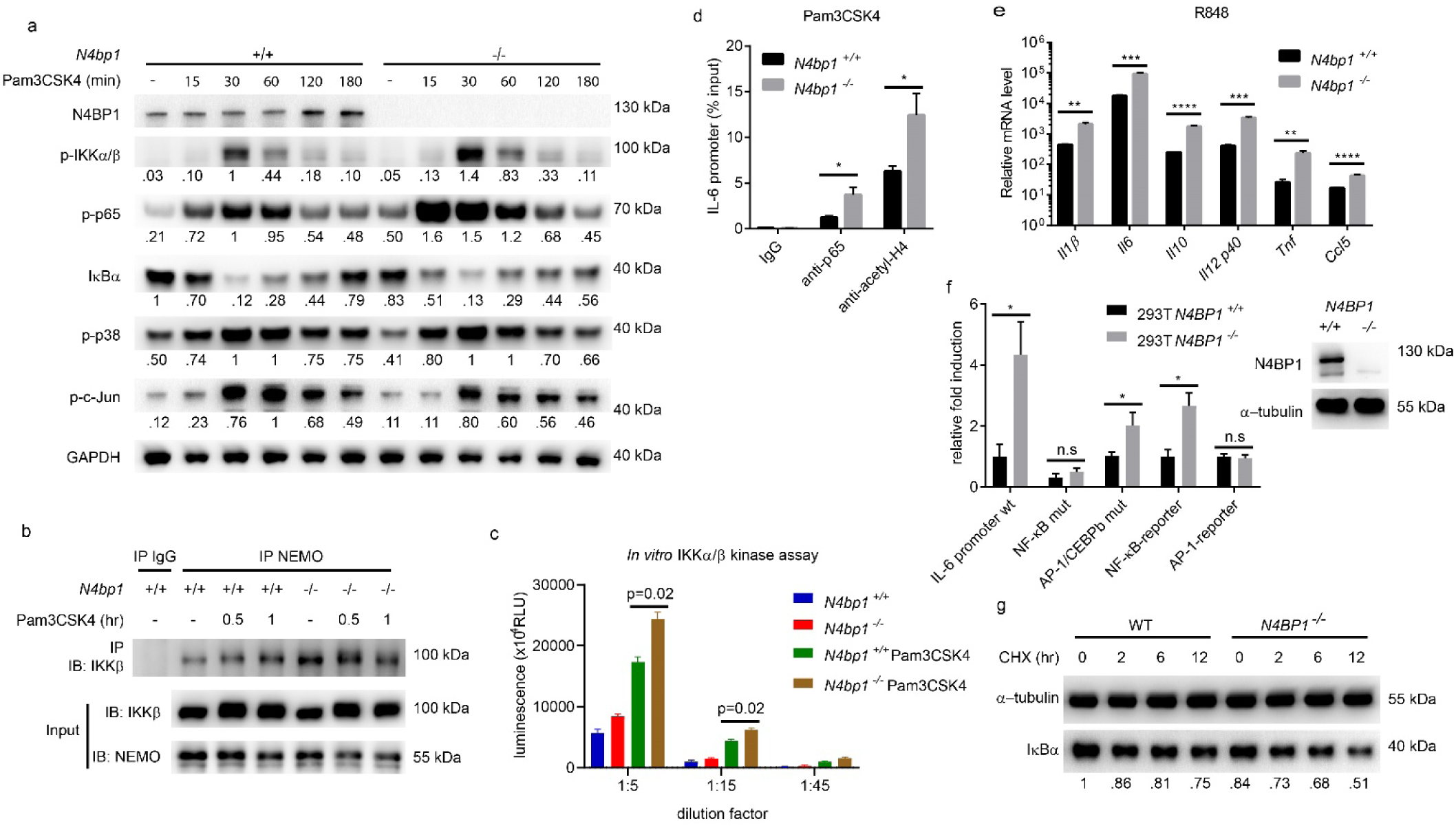
N4BP1 inhibits NF-κB activation and NF-κB-dependent gene expression. **a**, Immunoblot analysis of phosphorylated (p-) IKKα/β, p65, p38, and c-Jun in lysates of wild-type and *N4bp1*^*-/-*^ peritoneal macrophages stimulated with Pam3CSK4 (40 ng/ml) for the indicated times. Relative densitometric measurements are indicated below. **b**, Detection of the endogenous NEMO-IKKβ association in wild-type and *N4bp1*^*-/-*^ peritoneal macrophages stimulated with Pam3CSK4 for the indicated times, assessed by immunoprecipitation (IP) with rabbit IgG as a control, or with anti-NEMO, followed by immunoblot analysis with anti-IKKβ or anti-NEMO. **c**, *In vitro* IKKα/β kinase assay. The IKK complex was immunoprecipitated from *N4bp1*^*+/+*^ or *N4bp1*^*-/-*^ peritoneal macrophages with anti-NEMO. Cells were treated with or without Pam3CSK4 (40 ng/ml) for 30 min before collection. **d**, ChIP assay and qPCR of *Il6* promoter DNA in peritoneal macrophages 2 h after Pam3CSK4 stimulation. Cell lysates were immunoprecipitated with rabbit IgG, anti-p65, or anti-acetyl-histone H4. **e**, RT-qPCR analysis of *Il1β, Il6, Il10, Il12p40, Tnf*, and *Ccl5* in *N4bp1*^*+/+*^ and *N4bp1*^*-/-*^ peritoneal macrophages stimulated with R848. **f**, *Left*, Luciferase reporter activity dependent on the indicated promoters (X-axis) in wild-type or *N4BP1*^*-/-*^ HEK293T cells. *Right*, Immunoblot analysis of N4BP1 expression in wild-type and *N4BP1*^*-/-*^ HEK293T cells. **g**, Immunoblot analysis of IκBα in lysates of wild-type and *N4BP1*^*-/-*^ HEK293T cells treated with cycloheximide (CHX, 20 μg/ml) for the indicated times. Relative densitometric measurements of IκBα are indicated below. * *P* < 0.05, \*\**P* < 0.01, \*\*\**P* < 0.001, \*\*\*\**P* < 0.0001 (two-tailed Student’s *t*-test). Data are representative of two independent experiments (mean ± s.d. in **d**,**e**,**f**).

The amino acid sequence of N4BP1 is 81% identical in humans and mice. We knocked out endogenous N4BP1 via CRISPR-Cas9 gene targeting in the human monocytic cell line THP1 and observed more TNF secretion in response to stimulation with Pam3CSK4 and R848, but not LPS, relative to that of N4BP1-sufficient THP1 cells (Supplementary Fig. 1i-k). These data indicate that the function of N4BP1 in inhibiting TRIF-independent TLR signaling is conserved between humans and mice.

### N4BP1 inhibits NF-κB activation and NF-κB-dependent gene expression

We examined MAPK and NF-κB signaling in *N4bp1*^*-/-*^ peritoneal macrophages. We found that phosphorylated NF-κB p65 was markedly increased in *N4bp1*^*-/-*^ macrophages compared to *N4bp1*^*+/+*^ macrophages before or after Pam3CSK4 stimulation (Fig. 2a). Phosphorylated IKKα/β was also increased in *N4bp1*^*-/-*^ macrophages after Pam3CSK4 stimulation (Fig. 2a) and this correlated with increased IKKβ association with NEMO (Fig. 2b), an interaction required for NF-κB activation^3^. Indeed the IKKα/β kinase activity was significantly elevated in *N4bp1*^*-/-*^ macrophages (Fig. 2c). By contrast, we observed no appreciable difference between *N4bp1*^*-/-*^ and wild-type macrophages in their levels of phosphorylated p38 and c-Jun in response to Pam3CSK4 (Fig. 2a). Moreover, chromatin immunoprecipitation and quantitative PCR (ChIP-qPCR) indicated that greater amounts of IL-6 promoter DNA was associated with NF-κB p65 and acetyl-histone H4 in *N4bp1*^*-/-*^ macrophages than in wild-type macrophages stimulated with Pam3CSK4 (Fig. 2d). These findings demonstrate that NF-κB activation and recruitment to the IL-6 promoter was increased in *N4bp1*^*-/-*^ macrophages in response to Pam3CSK4.

The increased association between acetyl-histone H4 and the IL-6 promoter in *N4bp1*^*-/-*^ cells suggested that *Il6* was more transcriptionally active in *N4bp1*^*-/-*^ cells than in *N4bp1*^*+/+*^ cells. To investigate transcript expression of *Il6, Tnf*, and other genes, we performed RNA sequencing of *N4bp1*^*+/+*^ and *N4bp1*^*-/-*^ peritoneal macrophages before and after R848 or LPS stimulation. We found that the expression of NF-κB-dependent genes, including *Tnf* and *Il6*, was higher in *N4bp1*^*-/-*^ macrophages than in *N4bp1*^*+/+*^ macrophages after R848 but not LPS stimulation (Supplementary >Fig. 3a, b and Supplementary Data 1). These observations were confirmed by RT-qPCR of *Il1b, Il6, Il10, Il12 p40, Tnf*, and *Ccl5* (Fig. 2e). Moreover, we found that NF-κB-dependent genes were upregulated in unstimulated *N4bp1*^*-/-*^ macrophages relative to their expression levels in *N4bp1*^*+/+*^ macrophages (Supplementary Fig. 3c and Supplementary Data 1).

**Figure 3.**
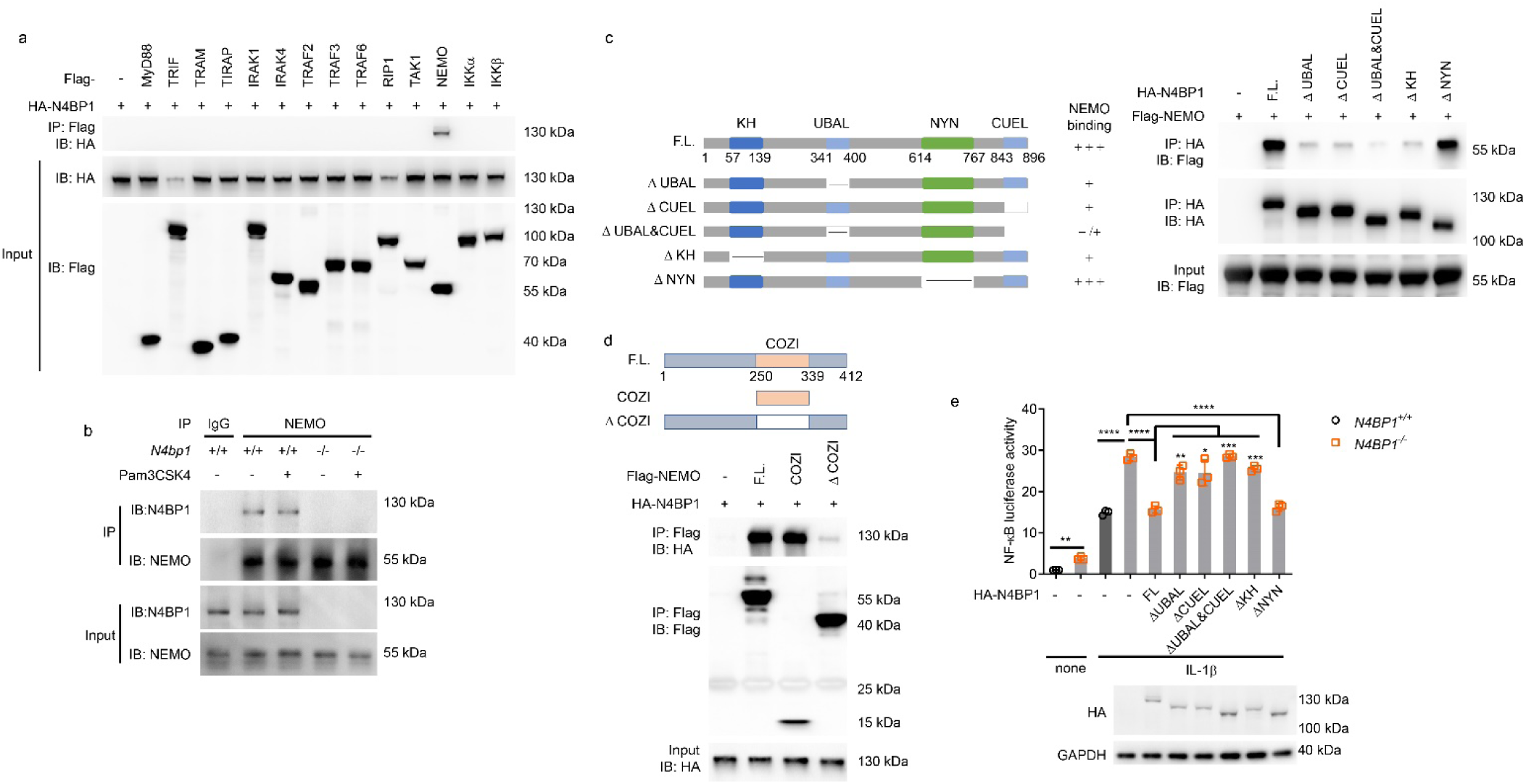
N4BP1 UBA-like and CUE-like domains associate with the NEMO COZI domain to inhibit NF-κB. **a**, Immunoprecipitation and immunoblot analysis of the interaction between hemagglutinin (HA)-tagged N4BP1 and Flag-tagged components of the NF-κB pathway in HEK293T cells transfected to overexpress HA-tagged N4BP1 (HA-N4BP1) alone (−) or together with vector encoding Flag-tagged NF-κB pathway components (above lanes). **b**, Detection of the endogenous N4BP1-NEMO association in wild-type and *N4bp1*^*-/-*^ peritoneal macrophages stimulated with Pam3CSK4 (+) or left unstimulated (−), assessed by immunoprecipitation with rabbit IgG as a control, or with anti-NEMO, followed by immunoblot analysis with anti-N4BP1 or anti-NEMO. **c**,**d**, Mapping the binding regions between HA-N4BP1 and Flag-NEMO. (**c**) *Left*, Schematic representation of N4BP1 deletion mutants and amount of NEMO they bind. *Right*, Immunoprecipitation and immunoblot analysis of the interaction between Flag-NEMO and HA-N4BP1 deletion mutants. (**d**) *Above*, Schematic representation of NEMO deletion mutants. *Below*, Immunoprecipitation and immunoblot analysis of the interaction between HA-N4BP1 and Flag-NEMO deletion mutants. **e**, *Above*, NF-κB-dependent luciferase reporter activity in wild-type or *N4BP1*^*-/-*^ HEK293T cells transfected with empty vector (-) or vectors encoding the indicated N4BP1 deletion mutants and stimulated with IL-1β or left unstimulated (none). Data points represent independent cultures. *Below*, Immunoblot analysis of HA-N4BP1 or truncation mutants in the pooled, analyzed cells. FL, full length N4BP1. * *P* < 0.05, \*\**P* < 0.01, \*\*\**P* < 0.001, \*\*\*\**P* < 0.0001 (two-tailed Student’s *t*-test). Data are representative of two independent experiments (mean ± s.d. in **e**).

To analyze the gating effect of N4BP1 on various promoter elements, we used CRISPR-Cas9 gene targeting to generate N4BP1 knockout HEK293T cells and transfected them with luciferase reporter constructs driven by (1) an IL-6 promoter, (2) an IL-6 promoter with the NF-κB binding site deleted, (3) an IL-6 promoter with AP-1 and CEBPb binding sites deleted^9^, (4) a pure NF-κB binding site, or (5) a pure AP-1 binding site. In the absence of stimulation, wild-type IL-6 promoter-dependent luciferase activity was elevated in *N4BP1*^*-/-*^ cells compared to that in *N4BP1*^*+/+*^ cells; this activity was abolished by deletion of the NF-κB binding site, but not deletion of the AP-1 and CEBPb binding sites, from the IL-6 promoter (Fig. 2f). Similarly, a consensus NF-κB binding site, but not an AP-1 binding site, drove increased reporter activity in *N4BP1*^*-/-*^ cells relative to *N4BP1*^*+/+*^ cells (Fig. 2f). Furthermore, the level of the NF-κB inhibitor IκBα was decreased in *N4BP1*^*-/-*^ HEK293T cells compared to that in *N4BP1*^*+/+*^ cells, both at steady state (t=0) and at various times after imposition of translational blockade with cycloheximide (Fig. 2g). Together, these data demonstrate that N4BP1 negatively regulates NF-κB-dependent gene expression in macrophages both before and in response to TLR stimulation.

### N4BP1 associates with NEMO to inhibit NF-κB

To identify the functional target(s) of N4BP1, we co-transfected different NF-κB pathway components, each together with N4BP1, into HEK293T cells. By immunoprecipitation (IP), NEMO was identified as a binding partner of N4BP1 (Fig. 3a). The interaction between endogenous N4BP1 and NEMO was also detected in peritoneal macrophages and was not influenced by cell stimulation with Pam3CSK4 (Fig. 3b).

N4BP1 was reported to have a K Homology (KH) domain, putatively capable of RNA binding, near the N-terminus, and a N4BP1, YacP-like nuclease (NYN) domain near the C-terminus^10^. To test whether N4BP1 directly degraded NF-κB-dependent gene mRNA, we treated wild-type and *N4BP1*^*-/-*^ HEK293T cells with IL-1β together with actinomycin D, an inhibitor of transcription. Increased IκBα and IL-8 mRNA levels were observed in *N4BP1*^*-/-*^ cells, however the decay rates were comparable in wild-type and *N4BP1*^*-/-*^ cells (Supplementary Fig. 4a, b). Furthermore, mice with N4BP1 NYN domain mutation Y774H did not affect R848 induced TNF production (Supplementary Fig. 1d). The results suggest that N4BP1 does not function as a nuclease to negatively regulate NF-κB-dependent gene expression.

**Figure 4.**
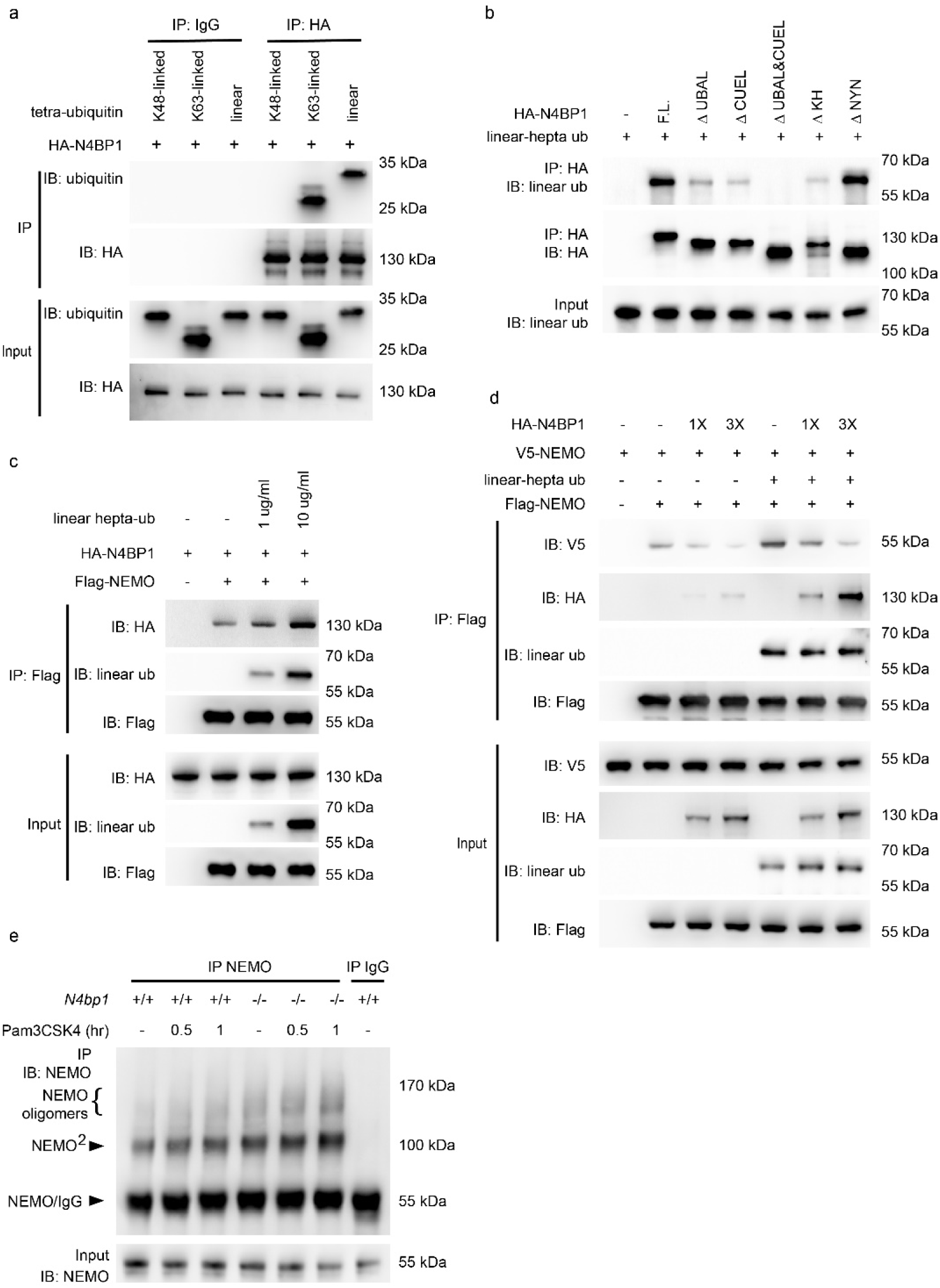
N4BP1 inhibits NEMO homo-oligomerization. **a**, Tetra-ubiquitin and purified recombinant HA-N4BP1 were incubated together and subjected to immunoprecipitation with mouse IgG or anti-HA. **b**, Linear hepta-ubiquitin was incubated without (-) or with purified recombinant HA-N4BP1 or its domain deletion forms and subjected to immunoprecipitation with anti-HA. **c**, Purified recombinant HA-N4BP1 and Flag-NEMO were incubated without (-) or with linear hepta-ubiquitin as indicated, and subjected to immunoprecipitation with anti-Flag. **d**, Purified recombinant Flag-NEMO and V5-NEMO were mixed together and then incubated with different amounts of HA-N4BP1 in the presence of linear hepta-ubiquitin or without ubiquitin, and subjected to immunoprecipitation with anti-Flag. **e**, Detection of the endogenous NEMO oligomerization in wild-type and *N4bp1*^*-/-*^ peritoneal macrophages, assessed by immunoprecipitation with rabbit IgG as a control, or with anti-NEMO, followed by immunoblot analysis with anti-NEMO. Cells were stimulated with Pam3CSK4 or left unstimulated (−) and treated with 2 mM disuccinimidyl suberate (DSS) for 10 min before collection. Complexes were analyzed by immunoblotting with the indicated antibodies (**a-e**). Data are representative of two independent experiments.

By searching COLIS^11^ and HHpred (https://toolkit.tuebingen.mpg.de/#/tools/hhpred), we further identified two putative ubiquitin binding domains in N4BP1: a ubiquitin-associated (UBA)-like domain in the middle region of N4BP1 (a.a. 341-400) and a ubiquitin conjugation to ER degradation (CUE)-like domain at the C-terminus (a.a. 843-896) (Fig. 3c). To map the region responsible for N4BP1-NEMO association, we generated constructs encoding HA-tagged N4BP1 deletion fragments (Fig. 3c), expressed them in HEK293T *N4BP1*^*-/-*^ cells, and examined the ability of the encoded proteins to bind to NEMO. The KH domain, UBA-like domain, and CUE-like domain were necessary for interaction with NEMO since deletion of any one of them reduced the association between N4BP1 and NEMO (Fig. 3c). The NYN domain in N4BP1 was dispensable for NEMO binding (Fig. 3c). In NEMO, the coil zipper (COZI) domain was necessary and sufficient for binding to N4BP1 (Fig. 3d).

We tested the effect of deletion of the UBA-like domain, CUE-like domain, or the KH domain of N4BP1 on NF-κB-dependent gene expression in *N4BP1*^*-/-*^ HEK293T cells. NF-κB driven luciferase reporter activity was increased in *N4BP1*^*-/-*^ HEK293T cells compared to that in *N4BP1*^*+/+*^ cells at steady state and after IL-1β stimulation (Fig. 3e). Transfection of *N4BP1*^*-/-*^ cells with either full-length N4BP1 or N4BP1 with NYN domain deletion reduced the luciferase activity to levels similar to those in wild-type HEK293T cells after IL-1β stimulation. N4BP1 with KH domain deletion or with one or both ubiquitin binding domains deleted failed to normalize NF-κB-dependent luciferase activity when expressed in *N4BP1*^*-/-*^ HEK293T cells (Fig. 3e). These findings indicate that the KH domain and ubiquitin binding domains of N4BP1 are necessary for its association with NEMO and the inhibition of NF-κB.

### N4BP1 inhibits NEMO oligomerization

We next explored the mechanism by which N4BP1 affected the function of NEMO. The UBA-like domain and CUE-like domain are potential sites for ubiquitin binding. Indeed, when incubated *in vitro*, purified recombinant HA-N4BP1 immunoprecipitated together with K63-linked and linear tetra-ubiquitin chains, but not with K48-linked tetra-ubiquitin chains (Fig. 4a). However, N4BP1 lacking the UBA-like and CUE-like domains (ΔUBAL&CUEL) failed to bind to linear and K63-linked ubiquitin chains (Fig. 4b and Supplementary Fig. 5a). NEMO ubiquitination and NEMO binding to linear ubiquitin chains are both required for NF-κB activation^12, 13^. We therefore hypothesized that ubiquitin chains might regulate the interaction between NEMO and N4BP1. We confirmed the direct interaction of purified recombinant Flag-NEMO and HA-N4BP1 proteins by immunoprecipitation (Fig. 4c and Supplementary Fig. 5b). The addition of purified recombinant polyubiquitin chains enhanced in a dose-dependent manner the interaction between NEMO and N4BP1 (Fig. 4c Supplementary Fig. 5b). These data indicate that linear and K63-linked ubiquitin chains, which bind to both N4BP1 and NEMO, promote N4BP1-NEMO interaction.

**Figure 5.**
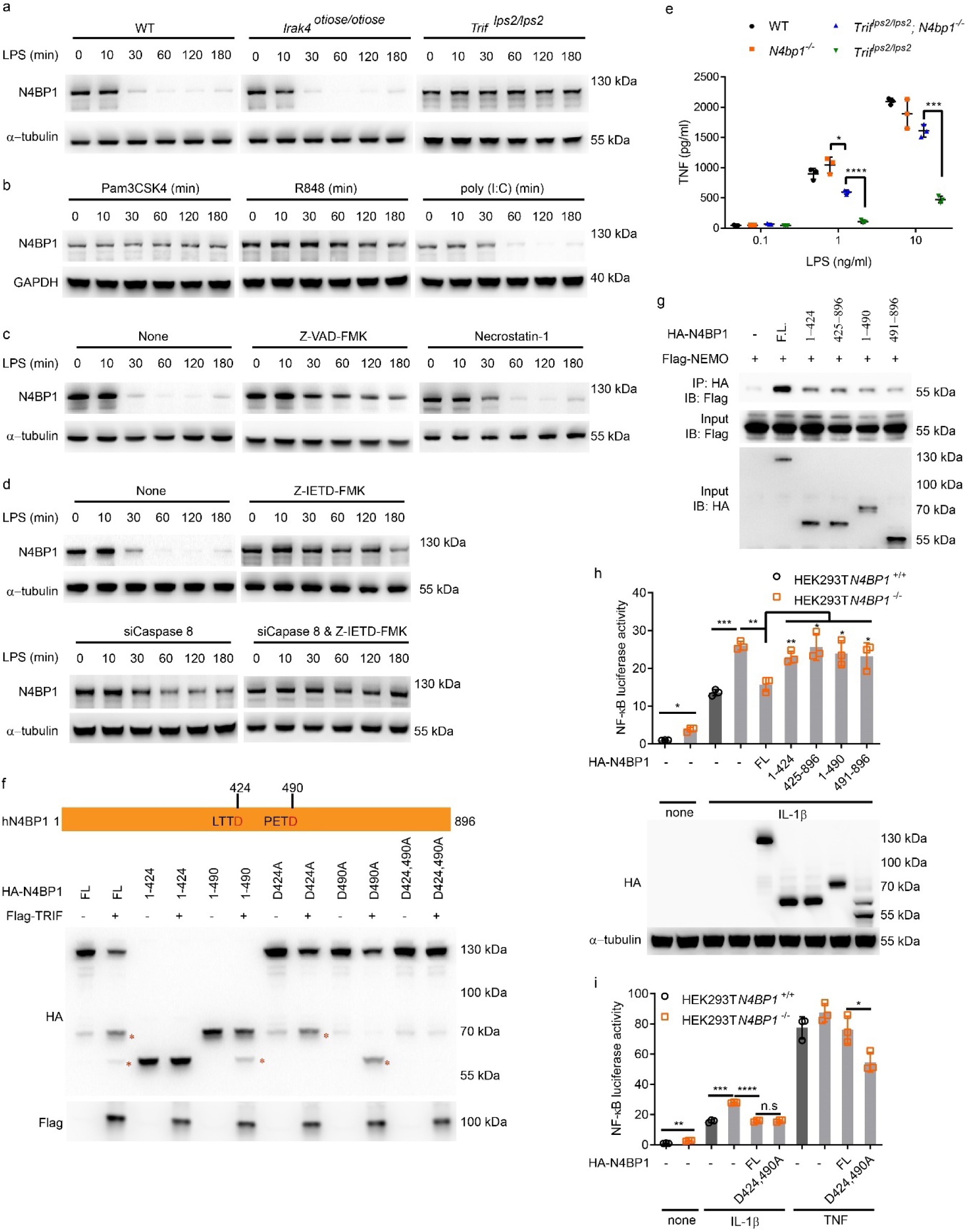
TRIF-dependent cleavage of N4BP1 by caspase 8. **a-d**, Immunoblot analysis of N4BP1 in wild-type, *Irak4*^*otiose/otiose*^, or *Trif*^*lps2/lps2*^ peritoneal macrophages stimulated with LPS for the indicated times (**a**), in wild-type peritoneal macrophages stimulated with Pam3CSK4 (40 ng/ml), R848 (20 ng/ml), or poly(I:C) for the indicated times (**b**), or in wild-type peritoneal macrophages pre-treated with Z-VAD-FMK(20 μM), Necrostatin-1 (50 μM), or Z-IETD-FMK (50 μM) for 1 h and then stimulated with LPS for the indicated times (**c**,**d**). **e**, TNF concentration in the culture medium of wild-type (WT), *N4bp1*^*-/-*^, *Trif*^*lps2/lps2*^*N4bp1*^*-/-*^, or *Trif*^*lps2/lps2*^ peritoneal macrophages treated with LPS at different concentrations (n=3 mice per genotype). **f**, Immunoblot analysis of HA-N4BP1 or truncation or point mutants of HA-N4BP1 expressed with or without TRIF in HEK293T cells. Asterisks indicate cleaved fragments of N4BP1. **g**, Detection of the interaction between HA-N4BP1 or truncation mutants of HA-N4BP1 and Flag-NEMO expressed in HEK293T cells, assessed by immunoprecipitation with anti-HA followed by immunoblot analysis with anti-Flag. **h**, *Above*, NF-κB-dependent luciferase reporter activity in wild-type or *N4BP1*^*-/-*^ HEK293T cells transfected with empty vector (-) or vectors encoding the indicated N4BP1 truncation mutants and stimulated with IL-1β or left unstimulated (none). *Below*, Immunoblot analysis of HA-N4BP1 or truncation mutants in the pooled, analyzed cells. **i**, NF-κB-dependent luciferase reporter activity in wild-type or *N4BP1*^*-/-*^ HEK293T cells transfected with empty vector (-) or a vector encoding the indicated N4BP1 mutant and stimulated with IL-1β, TNF, or left unstimulated (none). FL, full length N4BP1. Each symbol represents an individual mouse (**e**) or independent culture (**h**,**i**). * *P* < 0.05, \*\**P* < 0.01, \*\*\**P* < 0.001, \*\*\*\**P* < 0.0001 (two-tailed Student’s *t*-test). Data are representative of two independent experiments (mean ± s.d. in **e**,**h**,**i**).

Previous reports have shown that NEMO homo-oligomerization is required for recruitment of IKKα and IKKβ and subsequent NF-κB activation^14, 15^. In addition, such oligomerization may be mediated by a C-terminal region of NEMO that overlaps with the binding site for N4BP1^14, 16, 17^. Thus, we tested whether NEMO-N4BP1 interaction could inhibit NEMO homo-oligomerization. Purified recombinant V5-tagged NEMO immunoprecipitated with Flag-tagged NEMO and this interaction was enhanced in the presence of linear polyubiquitin chains (Fig. 4d). The addition of HA-tagged N4BP1 dose-dependently reduced the amount of V5-NEMO that immunoprecipitated with Flag-NEMO (Fig. 4d). Similar results were observed when K63-linked polyubiquitin chains were substituted for linear polyubiquitin chains in the same experiment (Supplementary Fig. 5c). We also observed more NEMO dimers and oligomers in *N4bp1*^*-/-*^ peritoneal macrophages (Fig. 4e). These findings suggest that N4BP1 blocks NEMO oligomerization by binding to the C-terminal oligomerization domain of NEMO, thereby precluding IKKα/β recruitment and NF-κB activation.

### TRIF-dependent N4BP1 cleavage by caspase 8

TLR proximal signaling pathways differ depending in part on the adapters that serve them, but all converge downstream from TRAF6 to signal activation of MAPKs and NF-κB^18^. To understand how N4BP1 deficiency specifically enhanced NF-κB signaling in response to Pam3CSK4 and R848, but had no effect on NF-κB activation in response to poly(I:C) or LPS, we first examined N4BP1 expression levels in peritoneal macrophages after LPS, Pam3CSK4, R848 and poly(I:C) stimulation. We found that N4BP1 expression decreased by 30 min after LPS treatment, was undetectable at 60 and 120 min after LPS treatment and recovered to barely detectable levels by 180 min after stimulation (Fig. 5a). The LPS-induced downregulation of N4BP1 was not observed in TRIF-deficient (*Trif*^*lps2/lps2*^) macrophages but remained intact in IRAK4-deficient macrophages homozygous for the *otiose* allele, which fail to propagate signals from MyD88 (Fig. 5a). Poly(I:C) stimulation of wild-type macrophages induced N4BP1 downregulation with a similar time course and level of reduction as that induced by LPS (Fig. 5b). In contrast, N4BP1 failed to be downregulated in wild-type macrophages in response to Pam3CSK4 and R848 (Fig. 5b). To summarize, LPS stimulation of wild-type or IRAK4-deficient macrophages, or poly(I:C) stimulation of wild-type macrophages, led to downregulation of N4BP1 expression. Downregulation of N4BP1 was not observed in TRIF-deficient macrophages treated with LPS or in wild-type macrophages treated with Pam3CSK4 or R848. These data indicate that stimulation of TRIF-dependent TLR signaling pathways results in N4BP1 downregulation.

We hypothesized that TRIF might direct the downregulation and inhibition of N4BP1. To investigate which pathway(s) might function downstream from TRIF to downregulate N4BP1, we tested whether inhibitors of the proteasome, autophagy, cysteine proteases, or protein translation could block LPS-induced N4BP1 downregulation, but none did so (Supplementary Fig. 6). Neither did the necrosis inhibitor Necrostatin-1 block LPS-induced N4BP1 downregulation (Fig. 5c). However, the pan-caspase inhibitor Z-VAD-FMK successfully blocked N4BP1 downregulation such that N4BP1 was detectable by immunoblot of wild-type macrophage lysates at all time points up to 180 min after LPS stimulation (Fig. 5c). We tested more specific caspase inhibitors and found that Z-IETD-FMK, a caspase 8 inhibitor, abrogated N4BP1 downregulation in response to LPS (Fig. 5d). Furthermore, we confirmed that caspase 8 knockdown inhibited LPS induced N4BP1 downregulation (Fig. 5d). These data suggest that LPS-induced TRIF-dependent signaling activates caspase 8 to cleave N4BP1, resulting in reduced levels of full length N4BP1. This is consistent with a previous report that TRIF but not MyD88 activates caspase 8 (19).

**Figure 6.**
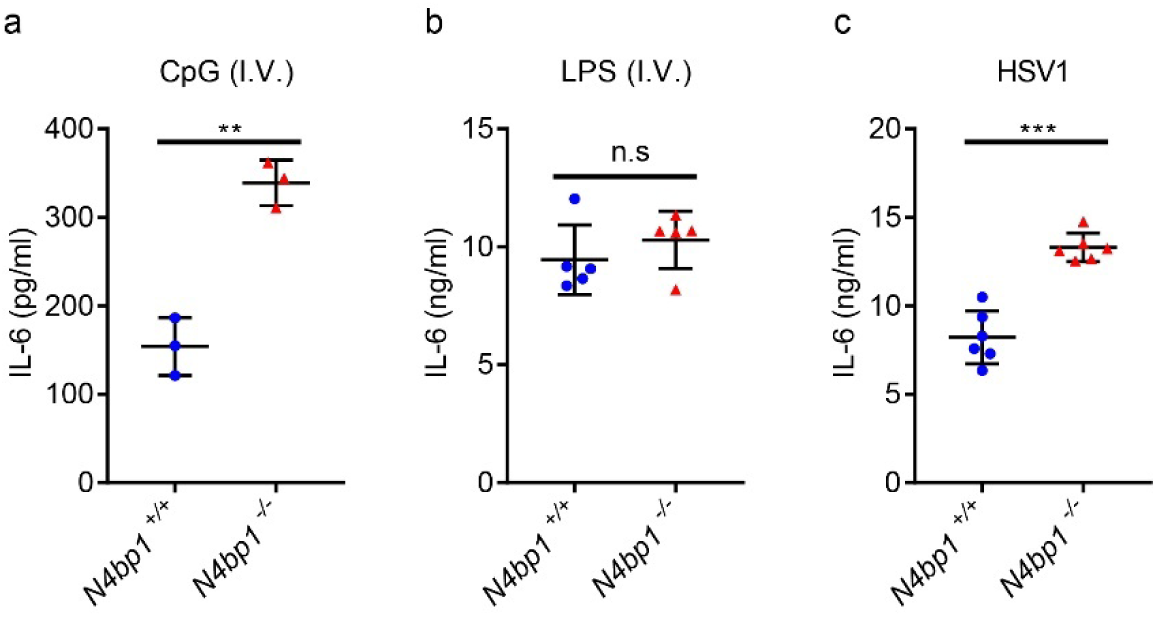
Enhanced response of *N4bp1*^*-/-*^ mice to *in vivo* CpG and HSV1 stimulations. **a-c**, Serum concentration of IL-6 in age matched wild-type and *N4bp1*^*-/-*^ mice intravenously (i.v.) injected with CpG (**a**), LPS (**b**), or HSV1 (**c**). Each symbol (**a-c**) represents an individual mouse. * *P* < 0.5, \*\**P* < 0.01, \*\*\**P*< 0.001 (two-tailed Student’s *t*-test). Data are representative of two independent experiments (mean ± s.d.).

To further probe the functional relationship between TRIF and N4BP1, we tested the effect of N4BP1 deficiency on LPS-induced TNF production by *Trif*^*lps2/lps2*^ macrophages. We observed increased TNF production by *Trif*^*lps2/lps2*^*N4bp1*^*-/-*^ cells compared to *Trif*^*lps2/lps2*^ cells, consistent with the hypothesis that TRIF signaling inhibits N4BP1 (Fig. 5e). As expected, wild-type and *Trif*^*lps2/lps2*^ macrophages produced similar amounts of TNF in response to Pam3CSK4 or R848 stimulation, and N4BP1 deficiency significantly increased those amounts (Supplementary Fig. 7a, b). Since TLR3 signaling is fully dependent on TRIF, no TNF production was detected in either *Trif*^*lps2/lps2*^*N4bp1*^*-/-*^ or *Trif*^*lps2/lps2*^ macrophages after poly(I:C) stimulation (Supplementary Fig. 7c). These data support the conclusion that TRIF-dependent signaling leads to inhibition of N4BP1.

TNF also induces caspase 8 activation^20^, and we indeed observed N4BP1 downregulation after we treated peritoneal macrophages with TNF but not with IL-1β (Supplementary Fig. 7d, e). In line with these findings, IL-1β but not TNF induced more *Il6* and *Tnf* mRNA expression in *N4bp1*^*-/-*^ macrophages than in wild-type macrophages (Supplementary Fig. 7f, g).

Caspase 8 preferentially cleaves protein and peptide substrates immediately after D residues in ‘XEXD’ and ‘LETD’ motifs, respectively, where ‘X’ represents any residue^21^. By scanning the human N4BP1 protein sequence, we identified two putative caspase 8 cleavage sites (LTTD424 and PETD490), which were also present in the mouse protein sequence. To investigate whether N4BP1 is cleaved at one or both sites in a TRIF-dependent manner, we expressed HA-tagged N4BP1 with or without Flag-tagged TRIF in HEK293 cells and immunoblotted the cell lysates for HA-N4BP1. In addition to the full-length protein, two N4BP1 fragments were detected in lysates of cells expressing HA-N4BP1 and Flag-TRIF, and these fragments corresponded in size to the 424-aa and 490-aa products expected to result from caspase 8 cleavage (Fig. 5f, lanes 1-6). A D424A mutation abrogated the 424-aa cleavage product while preserving the 490-aa product (Fig. 5f, lanes 7-8). An analogous outcome resulted from a D490A mutation (Fig. 5f, lanes 9-10). Mutation of both D424 and D490 to alanine blocked detectable TRIF-dependent N4BP1 cleavage (Fig. 5f, lanes 11-12). We also observed that HA-N4BP1 cleavage products immunoprecipitated reduced amounts of Flag-NEMO compared to full length HA-N4BP1 (Fig. 5g), and could not reduce NF-κB driven luciferase activity to a normal level when expressed in *N4BP1*^*-/-*^ HEK293T cells (Fig. 5h). When N4BP1 with D424A and D490A mutations was expressed in *N4BP1*^*-/-*^ HEK293T cells, it functioned the same as wild-type N4BP1 in reducing IL-1β induced NF-κB driven luciferase activity; in contrast, it diminished the luciferase response induced by TNF as compared to wild-type N4BP1 (Fig. 5i). Taken together, these results suggest that when activated during TRIF-dependent TLR signaling and TNF signaling, caspase 8 cleaves N4BP1 after D424 and/or D490. These cleavage events inactivate N4BP1 and reduce its binding to NEMO, resulting in unrestricted IKK complex formation and NF-κB activation.

### N4BP1 limits TRIF-independent NF-κB activation *in vivo*

To assess the role of N4BP1 as a checkpoint for TRIF-dependent and -independent TLR signaling *in vivo*, we intravenously injected wild-type and *N4bp1*^*-/-*^ mice with CpG or LPS and measured systemic cytokine responses. Three hours after CpG injection, the serum of *N4bp1*^*-/-*^ mice had a significantly higher concentration of IL-6 than did wild-type serum (Fig. 6a). However, we did not detect appreciable differences between *N4bp1*^*-/-*^ and wild-type mice in serum IL-6 concentration after LPS administration (Fig. 6b). These results were consistent with our previous observation that *N4bp1* deficiency enhanced TRIF-independent signaling.

The recognition of herpes simplex virus 1 (HSV1) via TLRs (notably TLR9) and cytosolic receptors^22^ initiates signaling leading to NF-κB activation. After infection with a non-lethal dose of HSV1, we monitored serum IL-6 in wild-type and *N4bp1*^*-/-*^ mice. Six hours after infection, *N4bp1*^*-/-*^ mice had significantly elevated serum concentrations of IL-6 compared to wild-type mice (Fig. 6c). Together, these results provide *in vivo* evidence that N4BP1 attenuates NF-κB activation.

## Discussion

N4BP1 was first identified as a NEDD4 binding protein^8^, and later as an inhibitor of the E3 ligase Itch, thereby stabilizing cell death regulators such as p73α and c-Jun^23^. N4BP1 interacts with K63-linked polyubiquitinated proteins including NEMO, A20, ABIN-1, and ABIN-2 (24). In 2011, N4BP1 was implicated as a negative regulator of NF-κB, although no mechanism was reported^25^. Most recently N4BP1 was shown to interact with the deubiquitinating enzyme CEZANNE^26^, which stabilizes TRAF3. When *N4BP1* was mutated in neuroblastoma cells, Spel et al. observed decreased TRAF3 expression resulting in enhanced canonical and noncanonical NF-κB signaling^26, 27, 28^. Furthermore, *Traf3* deficient cells produced elevated amounts of both TRIF-dependent and TRIF-independent proinflammatory cytokines^27, 28^. However, we found normal expression levels of TRAF3 in *N4bp1*^*-/-*^ cells and normal TNF and IL-6 responses by *N4bp1*^*-/-*^ macrophages to LPS and poly(I:C), raising the possibility that N4BP1 may fulfill distinct functions in macrophages vs. transformed neural cells, or that the effect of N4BP1 deficiency on NF-κB signaling in neuroblastoma cells may be independent of TRAF3.

Using unbiased phenotypic screening, we identified N4BP1 as an inhibitor of innate immune signaling mediated by TRIF-independent TLRs. Thus, N4BP1 deficiency resulted in excessive secretion of TNF and other NF-κB dependent cytokines in response to ligands for TLR2/TLR1, TLR7, and TLR9, but not ligands for TLR3 and TLR4. We demonstrated that the basis for this specificity is the TRIF-dependent activation of caspase 8, which cleaves and inactivates N4BP1. The synergistic effect of activation of TLR3 or TLR4 on cytokine production and NF-κB pathway activation induced by simultaneous stimulation of another TLR^29, 30, 31^ may be mediated in part by TRIF-initiated degradation of N4BP1. Extensive biochemical analyses demonstrated that N4BP1 inhibits TRIF-independent signaling through inhibition of NF-κB activation; N4BP1 binds to and blocks the oligomerization of NEMO that is necessary for activation of the IKK complex. Our finding that linear and K63-linked polyubiquitin chains enhanced the interaction between N4BP1 and NEMO demonstrates a previously unknown function for polyubiquitination in the regulation of NF-κB signaling.

The inactivation of N4BP1 specifically when TLR3 or TLR4 are engaged by their ligands (respectively dsRNA or LPS) implies a shared signaling need that is met by cleavage of N4BP1. While amplification of NF-κB signaling is one purpose of N4BP1 cleavage, another intriguing possibility involves the NYN domain (aa 491-896), characterized by a protein fold common to RNases of the PIN (PilT N-terminal) and FLAP/5’-3’ exonuclease superfamilies^32^. A PIN-like functional RNase domain is found in the innate immune regulator Zc3h12a, where it is important for IL-6 and IL-12 p40 mRNA decay following TLR stimulation^33^. Because the cleavage of N4BP1 by caspase 8 would leave the NYN domain intact, it is possible that the putative RNase activity of the NYN domain-containing fragment contributes to antiviral defense, acting in conjunction with the type I IFN response induced by TLR3, consistent with a recent report^34^. It has previously been suggested that the ancestral ligand specificity of TLR4 may have been for viral proteins^35^, since TLR4, like TLR3, is capable of recruiting and activating TRIF and therefore type I interferon responses (best suited to antiviral defense rather than antibacterial defense); TLR4 retains the ability to recognize certain viral proteins^36^. Immune responses to viruses detected by TRIF-independent TLRs, in contrast, appear to benefit from N4BP1 deficiency. We found that mice lacking N4BP1 were more capable than their wild-type littermates of coping with at least one such viral challenge (HSV1). Insofar as they show exaggerated innate immune responses but are not overtly suffering from autoimmunity or inflammation disease, pharmacologic inhibition of N4BP1 might have useful applications, for example in the improvement of vaccine responses or as an adjunct to checkpoint inhibitors in the treatment of neoplastic diseases.

In addition to TRIF-dependent TLR signaling, we also found that TNF signaling overcomes N4BP1-mediated inhibition of NF-κB. These pathways have in common the activation of caspase 8, suggesting that caspase 8 is a general switch controlling the strength of NF-κB signaling. These data also suggest that other pathways that activate caspase 8 may amplify NF-κB-dependent gene expression by inactivating N4BP1. MALT1 was also reported to cleave N4BP1 in T cells, suggesting that distinct proteases may be induced to cleave N4BP1 in a cell type- and/or ligand-specific manner^34^. In view of the requirement for caspase 8 for normal T cell and B cell proliferation and homeostasis in conditional knockout mice^37, 38^ and our finding that N4BP1 is also similarly required, we conclude that in some cases the two proteins must function independently, or that the cleavage of N4BP1 promotes event(s) that are needed for lymphocyte homeostasis. Mice lacking N4BP1 have fewer T cells in the peripheral blood, affecting both CD4^+^ and CD8^+^ cells, especially those with a naïve phenotype. While T lymphopenia has not been studied in detail, it might result from activation-induced cell death (presumably T cell-intrinsic), or as a compensatory response to the altered cytokine environment (presumably T cell-extrinsic). Adoptive transfer studies might be used to distinguish between these possibilities.

While this manuscript was in revision, Gitlin et al. reported similar findings showing that N4BP1 negatively regulates cytokine responses induced by TRIF-independent TLRs and that N4BP1 cleavage by caspase 8 contributes to the licensing effect of TNF on cytokine responses to TRIF-independent TLR activation^39^. Beyond this, we show the mechanism by which N4BP1 negatively regulates cytokine responses involves binding and inhibiting NEMO oligomerization, consequently preventing NF-κB activation.

## Methods

### Mice

Eight-to twelve-week-old male and female mice (Mus musculus) on a pure C57BL/6J background were used in experiments. Male C57BL/6J mice purchased from The Jackson Laboratory were mutagenized with N-ethyl-N-nitrosourea as described^40^. Mutagenized G0 males were bred to C57BL/6J females, and the resulting G1 males were crossed to C57BL/6J females to produce G2 mice. G2 females were backcrossed to their G1 sires to yield G3 mice, which were screened for phenotype. Whole-exome sequencing and mapping were performed as described^41^. *N4bp1*^*acorn*^ (https://mutagenetix.utsouthwestern.edu/phenotypic/phenotypic_rec.cfm?pk=3555), *N4bp1*^*stash*^ (https://mutagenetix.utsouthwestern.edu/phenotypic/phenotypic_rec.cfm?pk=5124), *N4bp1*^*walnut*^ (https://mutagenetix.utsouthwestern.edu/phenotypic/phenotypic_rec.cfm?pk=3890), *N4bp1*^*winter*^ (https://mutagenetix.utsouthwestern.edu/phenotypic/phenotypic_rec.cfm?pk=3787) and *Ticam1*^*lps2*^ (*Trif*^*lps2*^) (https://mutagenetix.utsouthwestern.edu/phenotypic/phenotypic_rec.cfm?pk=90) mutant strains were generated by mutagenesis with N-ethyl-N-nitrosourea and are described online.

All experimental procedures using mice were approved by the Institutional Animal Care and Use Committee of the University of Texas Southwestern Medical Center and were conducted in accordance with institutionally approved protocols and guidelines for animal care and use. All mice were maintained at the University of Texas Southwestern Medical Center in accordance with institutionally approved protocols.

### Generation of *N4bp1*^*-/-*^ mice and cells

CRISPR-Cas9-mediated gene targeting system was used to generate *N4bp1* knockout (KO) alleles. Female C57BL/6J mice were superovulated by injection of 6.5 U pregnant mare serum gonadotropin (Millipore), followed by injection of 6.5 U human chorionic gonadotropin (Sigma-Aldrich) 48 h later. The superovulated mice were subsequently mated overnight with C57BL/6J male mice. The following day, fertilized eggs were collected from the oviducts and in vitro-transcribed Cas9 mRNA (50 ng/μl) and N4bp1 small base-pairing guide RNA (50 ng/μl; 5’-TATCAAGGGGATCTGCGAGC-3’) were injected into the cytoplasm or pronucleus of the embryos. The injected embryos were cultured in M16 medium (Sigma-Aldrich) at 37 °C in 95% air plus 5% CO2. To produce mutant mice, two-cell stage embryos were transferred into the ampulla of the oviduct (10–20 embryos per oviduct) of pseudopregnant Hsd:ICR (CD-1) female mice (Harlan Laboratories). Chimeric mutant mice were first crossed with C57BL/6J mice and their offspring were intercrossed for the generation of *N4bp1*^*-/-*^ mice. *N4bp1*^*-/-*^ mice have a four–base pair deletion (italics) in *N4bp1* exon 2 (AGATCCCCTT *GCGA* GCCGGAGCTGGA). The *N4bp1*^*-/-*^ mice were genotyped by capillary sequencing with 5’-CTGGTGAATTGGTCTAACTTTGTC-3’ and 5’-AAACTGCTGAATGTGACTCC-3’ as the PCR primers and 5’-AAACTGCTGAATGTGACTCC-3’ as the sequencing primer.

For generation of *N4BP1*^*-/-*^ HEK293T or THP1 cells, cells were infected with lentivirus (lentiCRISPR v2) encoding sgRNA 5’-CACGCCTTGTTCTCCACTGA-3’ and Cas9. After puromycin selection, single cells were selected for subculture and immunoblotting was used to confirm knockout of N4BP1 expression. For reconstitution of *N4BP1*^*-/-*^ cells with full-length N4BP1 or its truncation mutants, lentiviral vector pBOBI cs2.0 N-HA harboring CRISPR-resistant sequence 5’-CTGTATTTGTgCCaCAaTGGAG-3’ (without changing amino acids) was used.

### Reagents

Ultra-pure LPS, Pam3CSK4, poly(I:C), R848, and IM-54 were obtained from Enzo Life Sciences. Z-IETD-FMK, recombinant mouse TNF, IL-1β, IFNγ, and human IL-1β were from R&D Systems. Disuccinimidyl suberate (DSS) was from Pierce. Cycloheximide (CHX), Z-VAD-FMK, Caspase-3/7 Inhibitor I, Actinomycin D, Bafilomycin A1, E-64 and 3-Methyladenine (3-MA) were from Sigma. Necrostatin-1 was from Abcam. MG132 was from EMD Millipore. cDNAs encoding N4BP1 and NEMO were amplified by standard PCR techniques and were subsequently inserted into mammalian expression vectors. All point mutations were introduced with a QuickChange II XL site-directed mutagenesis kit (Agilent Technologies). All constructs were confirmed by sequencing. Recombinant K48-linked, K63-linked and linear tetra-ubiquitin were obtained from LifeSensors. Caspase 8 siRNA was from Dharmacon. Antibodies to the following were used: mouse N4BP1 (EPNCIR118), human N4BP1 (ab209103) and NEMO (EPR16629) (Abcam); HA (6E2, HRP conjugate), GAPDH (8884), p65 (D14E12), p-p65 (93H1), p-IKKα/β (16A6), IκBα(L35A5), p-p38 (4511), p-c-Jun (D47G9), Histone H3 (9715), and GAPDH (D16H11) (Cell Signaling Technology); Flag (M2), V5 (V8012), β-actin (A2228) and α-tubulin (T6199) (Sigma-Aldrich); Acetyl-Histone H4 (PA5-40084, Invitrogen); Linear ubiquitin (LUB9, Millipore). Primary antibodies used for flow cytometry were: CD3ε (clone 145-2C11), CD4 (clone RM4-5), CD8α (clone 53-6.7), B220 (CD45R, clone RA3-6B2), NK-1.1 (clone PK136), CD44 (clone IM7), CD62L (clone MEL-14), CD11c (clone HL3), CD11b(clone M1/70), F4/80 (clone BM8), CD16/32 (Clone 2.4G2) (BD Biosciences).

None of the cell lines used (THP-1 and HEK293T) are listed in the database of commonly misidentified cell lines maintained by the International Cell Line Authentication Committee and National Center for Biotechnology Information BioSample.

### Isolation and culture of peritoneal macrophages, bone marrow–derived macrophages, and bone marrow–derived dendritic cells

Macrophages were elicited by intraperitoneal injection of 2 ml BBL thioglycollate medium, brewer modified (4%; BD Biosciences), and recovered 4 d later by peritoneal lavage with 5 ml phosphate-buffered saline (PBS). The peritoneal macrophages were cultured at 37 °C in 95% air and 5% CO2 in DMEM cell culture medium (DMEM containing 10% FBS (BioFluid), 1% penicillin and streptomycin (Life Technologies)). Mouse bone marrow–derived macrophages were collected by flushing of bone marrow cells from femurs and tibiae of mice. These cells were cultured for 7 d in DMEM cell culture medium containing 40 ng/ml macrophage colony-stimulating factor (M-CSF) (PeproTech). For the generation of bone marrow-derived dendritic cells, bone morrow cells were cultured in Petri dishes in 10 ml DMEM cell culture medium containing 20 ng/ml of mouse granulocyte-macrophage colony-stimulating factor (GM-CSF) (R&D Systems). On day 3 of culture, this was replaced with fresh medium containing GM-CSF. Loosely adherent cells were transferred to a fresh Petri dish and cultured for an additional 4 d.

### Measurement of cytokine production

Cells were seeded onto 96-well plates at a density of 1 ×10^5^ cells per well and then were stimulated as follows: Pam3CSK4 (40 ng/ml), R848 (20 ng/ml), LPS (10 ng/ml), or poly(I:C) (100 μg/ml) for 4 h or as indicated. THP1 cells were stimulated with Pam3CSK4 (100 ng/ml), R848 (100 ng/ml), and LPS (10 ng/ml) for 4 h. Cytokine concentrations in the supernatants were measured with ELISA kits for human TNF, mouse IL-6 and TNF (eBioscience).

### Immunoblotting

Cells or tissues (with homogenization) were lysed in RIPA buffer (50 mM Tris-Cl, pH 7.5, 150 mM NaCl, 1 mM EDTA, 1 mM EGTA, 1% vol/vol NP-40, 0.5% wt/vol sodium deoxycholate, 0.1% wt/vol SDS, 1mM dithiothreitol, plus Halt™ Protease and Phosphatase inhibitor Cocktail (Thermo Scientific)) immediately before use, and protein contents were determined by BCA assay (Thermo Scientific). In some cases, cells were directly lysed in 1X SDS sample buffer (50 mM Tris-Cl, pH 6.8, 2% wt/vol SDS, 5% vol/vol β-mercaptoethanol, 0.1% wt/vol bromophenol blue, and 10% vol/vol glycerol). Equal amounts (∼20 μg) of protein extracts were separated by electrophoresis on 4-12% NuPAGE Bis-Tris Mini Gels (Invitrogen) and then transferred to nitrocellulose membranes (Bio-Rad). The membrane was blocked for 1 h in Tris-buffered saline containing 0.1% vol/vol Tween 20 and 5% wt/vol non-fat milk and then probed with various primary antibodies overnight, followed by secondary antibodies conjugated to horseradish peroxidase. The immunoreactivity was detected with SuperSignal West Dura Chemiluminescent Substrate (Thermo Scientific).

### Immunoprecipitation

Cells were lysed in cold NP-40 lysis buffer (1% vol/vol NP-40, 50 mM Tris-HCl, pH 7.4, and 150 mM NaCl) supplemented with Halt™ Protease and Phosphatase inhibitor Cocktail. HA-tagged proteins were immunoprecipitated with anti-HA and ChIP-Grade Protein G Magnetic Beads (Cell Signaling Technology). Flag-tagged proteins were immunoprecipitated with Anti-FLAG M2 Magnetic Beads (Sigma-Aldrich). For the endogenous interaction assay, macrophages were lysed with NP-40 lysis buffer with protease inhibitors. The cell lysates were incubated overnight at 4 °C with NEMO antibody and ChIP-Grade Protein G Magnetic Beads (Cell Signaling Technology).

### Chromatin immunoprecipitation

Cells were stimulated with Pam3CSK4 (40 ng/ml) for 1 h. ChIP was conducted using SimpleChIP kit (Cell Signaling Technology) according to the manufacturer’s protocol. IL-6 promoter primers were the following: 5’-AATGTGGGATTTTCCCATGA-3’ (Forward) and 5’-GCTCCAGAGCAGAATGAGCTA-3’ (Reverse).ChIP samples were analyzed by real-time qPCR with the SYBR-Green Master Mix system (Life Technologies).

### Reporter analysis

Wild-type or *N4BP1*^*-/-*^ HEK293T cells seeded on 24-well plates were transiently transfected with 100 ng of the luciferase reporter plasmid together with a total of 800 ng of various expression plasmids or empty control plasmids. As an internal control, 10 ng of pRL-SV40 was transfected simultaneously. 24 h after transfection, the cells were stimulated with IL-1β (50 ng/ml) for 8 h. Dual luciferase activity in the total cell lysates was quantified (Promega).

### Quantitative real-time PCR

Total RNA was isolated from cells using the RNeasy RNA extraction kit (Qiagen), and cDNA synthesis was performed using 1 μg of total RNA (iScript, Bio-Rad). qPCR was performed with the following gene-specific primers: *Il1β*, 5’-TGTAATGAAAGACGGCACACC-3’ (fwd), 5’-TCTTCTTTGGGTATTGCTTGG-3’ (rev); *Il6*, 5’-CTCTGCAAGAGACTTCCATCC-3’ (fwd), 5’-CGACTTGTGAAGTGGTATAGACAG-3’ (rev); *Il10*, 5’-TGGCCCAGAAATCAAGGAGC-3’ (fwd), 5’-CAGCAGACTCAATACACACT-3’ (rev); *Il12p40*, 5’-GGAAGCACGGCAGCAGAATA-3’ (fwd), 5’-AACTTGAGGGAGAAGTAGGAATGG-3’ (rev); *Tnf*, 5’-CTGTAGCCCACGTCGTAGC-3’ (fwd), 5’-TTGAGATCCATGCCGTTG-3’ (rev); *Ccl5*, 5’-ACTCCCTGCTGCTTTGCCTAC-3’ (fwd), 5’-TGCTGCTGGTGTAGAAATACT-3’ (rev); *Gapdh*, 5’-CGTCCCGTAGACAAAATGGT-3’ (fwd), 5’-TTGATGGCAACAATCTCCAC-3’ (rev); *huGAPDH*, 5’-ACCCACTCCTCCACCTTTGA-3’ (fwd), 5’-CTGTTGCTGTAGCCAAATTCGT-3’ (rev); *huNfkbia* (*IκBα)*, 5’-CTCCGAGACTTTCGAGGAAATAC-3’ (fwd), 5’-GCCATTGTAGTTGGTAGCCTTCA-3’ (rev); *huIL8*,5’-ATAAAGACATACTCCAAACCTTTCCAC-3’ (fwd), 5’-AAGCTTTACAATAATTTCTGTGTTGGC-3’(rev).

### *In vitro* binding assay

Tagged NEMO and N4BP1 and its domain deletion forms were overexpressed in HEK293T cells. Fresh lysates from these HEK293T cells were prepared with RIPA lysis buffer and sonication. Soluble lysates were immunoprecipitated and eluted with Flag, HA or V5 peptides (Sigma). The eluate was run on a gel-filtration chromatography column (Superdex 200 Increase 10/300 GL; GE Healthcare) in assay buffer (50 mM Tris-HCl, pH 7.4, 150 mM NaCl and 1mM dithiothreitol). Peak fractions were concentrated with Amicon Ultra 10K filtration units (Millipore). The recombinant proteins were incubated with or without K63-linked, or linear tetra or hepta-ubiquitin in assay buffer for 2 h at 4 °C.

### *In vivo* challenge with CpG and LPS

Age- and sex-matched mice were challenged by intravenous injection of CpG (2 μg premixed with 15 μl DOTAP (Roche) in DPBS) or LPS (2 μg per mouse). Two (for LPS) or three hours (for CpG) after the injection, mice were bled from the retro-orbital sinus, and the concentration of IL-6 in the serum was assayed by ELISA.

### HSV-1 Infection

Age-matched mice were infected with HSV-1 (strain KOS, prepared and provided by the laboratory of Z.J. Chen, University of Texas Southwestern Medical Center, Dallas, TX) at 10^6^ PFU per mouse via intravenous injection. IL-6 in the serum was assayed 6 h after infection by ELISA. Body weight of the infected mice was monitored daily for 9 d.

### Flow cytometry

Equal amounts of blood were collected in Minicollect Tubes (Mercedes Medical) and centrifuged at 700g for separation of serum, and red blood cells remaining in the serum were lysed using RBC lysis buffer (eBioscience) before staining of immune cells and flow cytometry analysis. Cells were incubated with monoclonal antibody to CD16/32 and were labeled for 1 h at 4 °C using fluorochrome-conjugated monoclonal antibodies to mouse CD3ε, CD4, CD8α, CD44, CD62L, B220, NK1.1, CD11b, F4/80 and CD11c. Counting beads (Invitrogen) were added to the samples before collecting data. Data were acquired using an LSRFortessa Cell Analyzer (BD Biosciences).

### Nuclear-cytoplasmic fractionation

Peritoneal macrophages were stimulated with Pam3CSK4 for the indicated time periods. Nuclear-cytoplasmic fractionation was conducted using the NE-PER Nuclear and Cytoplasmic Extraction Reagents kit (Thermo Fisher Scientific) according to the manufacturer’s protocol.

### *In vitro* IKK kinase assay

Peritoneal macrophages were untreated or stimulated with Pam3CSK4 for 30 minutes. Cells were collected and lysed in cold NP-40 lysis buffer (1% vol/vol NP-40, 50 mM Tris-HCl, pH 7.4, and 150 mM NaCl) supplemented with Halt™ Protease and Phosphatase inhibitor Cocktail. The same amount of protein was immunoprecipitated with anti-NEMO and ChIP-Grade Protein G Magnetic Beads (Cell Signaling Technology). The IKKα/β kinase activity in the IP complex was tested by using IKKβ Kinase Enzyme Kit (Promega) and following the manufacturer’s instructions.

### Statistical Analysis

Comparisons of differences were between two unpaired experimental groups in all cases. An unpaired *t*-test (Student’s *t*-test) is appropriate and was used for such comparisons. One-way ANOVA was applied to experiments with three or more than three groups. The phenotypic performance of mice (C57BL/6J) and primary cells of these mice is expected to follow a normal distribution, as has been observed in large data sets from numerous phenotypic screens conducted by our group. Variation within each data set obtained by measurements from mice or primary cells was assumed to be similar between genotypes since all strains were generated and maintained on the same pure inbred background (C57BL/6J); experimental assessment of variance was not performed.

The statistical significance of differences between experimental groups was determined with GraphPad Prism 7 software and the Student’s *t* test (unpaired, two-tailed). A *P* value of <0.05 was considered statistically significant. * *P* < 0.05; ** *P <* 0.01; *** *P <* 0.001; **** *P <* 0.0001. No pre-specified effect size was assumed, and in general three mice or more for each genotype or condition were used in experiments; this sample size was sufficient to demonstrate statistically significant differences in comparisons between two unpaired experimental groups by an unpaired *t*-test. The investigator was not blinded to genotypes or group allocations during any experiment.

## Acknowledgements

We thank Hong Zhang (University of Texas Southwestern Medical Center) for helpful suggestions. This work was supported by the National Institutes of Health grants R01 AI125581 and U19 AI100627 (to B.B.) and by the Lyda Hill Foundation. H.S., Y.W., X.Z., X.L., M.T., P.A., S.H., J.Q., S.L., and B.B. received salary support from Pfizer, Inc.

## Author Contributions

H.S. and B.B. designed research; H.S., L.S., Y.W., A.L., X.Z., X.L., M.T., P.A., J.Q., and S.L. performed research; S.H. performed computational analysis; H.S. and B.B. analyzed data; and H.S., E.M.Y.M., and B.B. wrote the manuscript.

