## Supplementary Information for "Mechanism of negative regulation of NF-κB by N4BP1"

Supplementary Figure 1

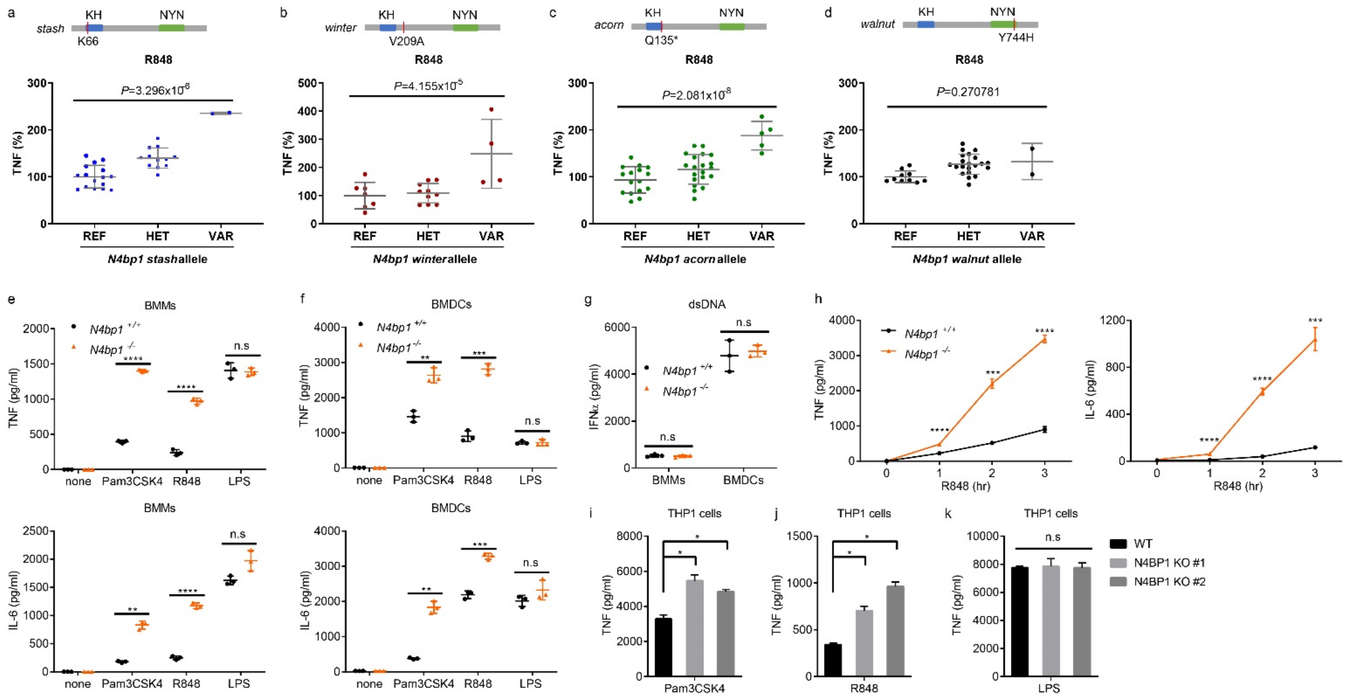


**Supplementary Figure 1. Enhanced TRIF-independent TNF and IL-6 production by N4BP1-deficient mouse BMMs, BMDCs and human monocytes. a-d,** Mutants in Fig.1b were shown individually with each p value. (**a**) *stash*, (**b**) *winter*, (**c**) *acorn*, (**d**) *walnut.* **e,f,** TNF (above) or IL-6 (below) concentration in the culture medium of *N4bp1*$^{+/+}$ and *N4bp1*$^{-/-}$ BMMs (**e**) or BMDCs (**f**) treated with the indicated stimuli. n=3 mice per genotype. **g,** IFNα concentration in the culture medium of *N4bp1*$^{+/+}$ and *N4bp1*$^{-/-}$ BMMs and BMDCs treated with dsDNA. n=3 mice per genotype. **h,** Time course analysis of TNF (above) or IL-6 (below) concentration in the culture medium of *N4bp1*$^{+/+}$ and *N4bp1*$^{-/-}$ peritoneal macrophages treated with R848. n=4 mice per genotype. **i-k,** TNF concentration in the culture medium of wild-type or *N4BP1*$^{-/-}$ THP1 cells treated with Pam3CSK4 (**i**), R848 (**j**) or LPS (**k**). Each symbol (**a-g**) represents an individual mouse. * $P < 0.05$, **$P < 0.01$, ***$P < 0.001$, ****$P < 0.0001$ (two-tailed Student's *t*-test). Data are representative of two independent experiments (mean ± s.d.) **(e-k)**.

Supplementary Figure 2

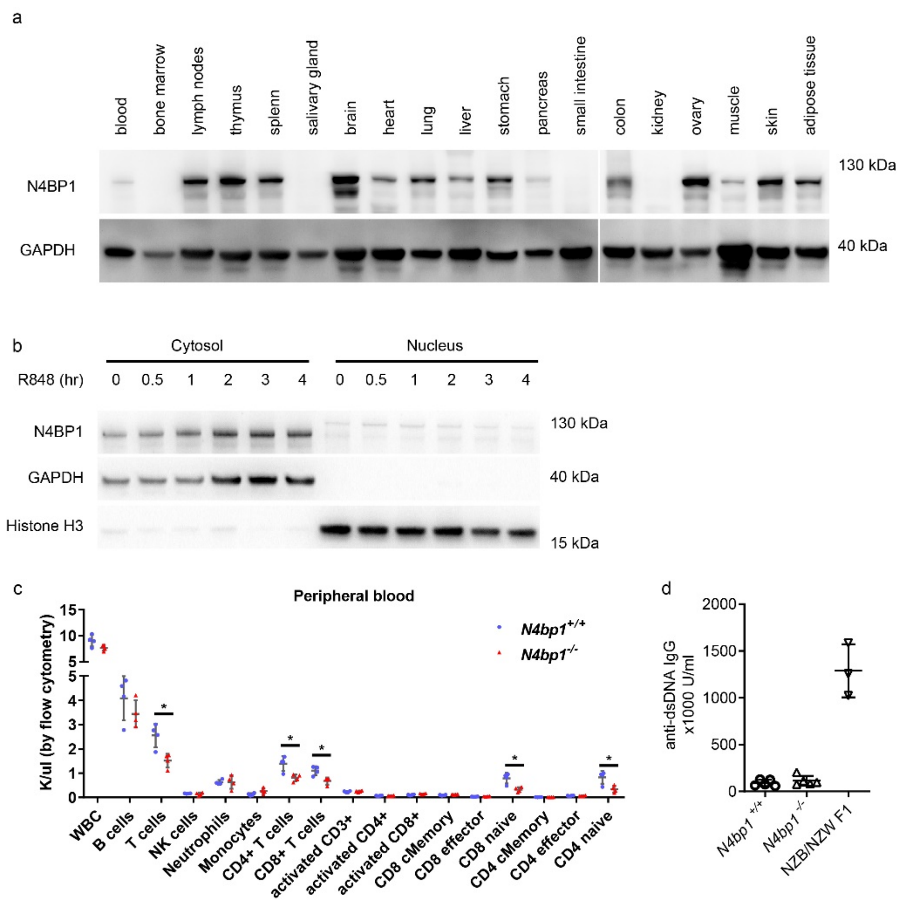


**Supplementary Figure 2. N4BP1-expressing tissues, and immune cell populations in peripheral blood and dsDNA antibodies in *N4bp1*$^{-/-}$ mice. a, b,** Immunoblot analysis of N4BP1 expression in different tissues (**a**) or cellular compartments of peritoneal macrophages (**b**). **c,** Quantification of lymphoid and myeloid cells analyzed by flow cytometry in the peripheral blood of wild-type and *N4bp1*$^{-/-}$ mice. **d,** Serum concentration of anti-dsDNA in wild-type and *N4bp1*$^{-/-}$ mice. NZB/NZW F1 mice were used as positive controls. Each symbol (**c,d**) represents an individual mouse. * $P < 0.05$ (two-tailed Student's *t*-test). Data are representative of two independent experiments (mean ± s.d. in **c,d**).

Supplementary Figure 3

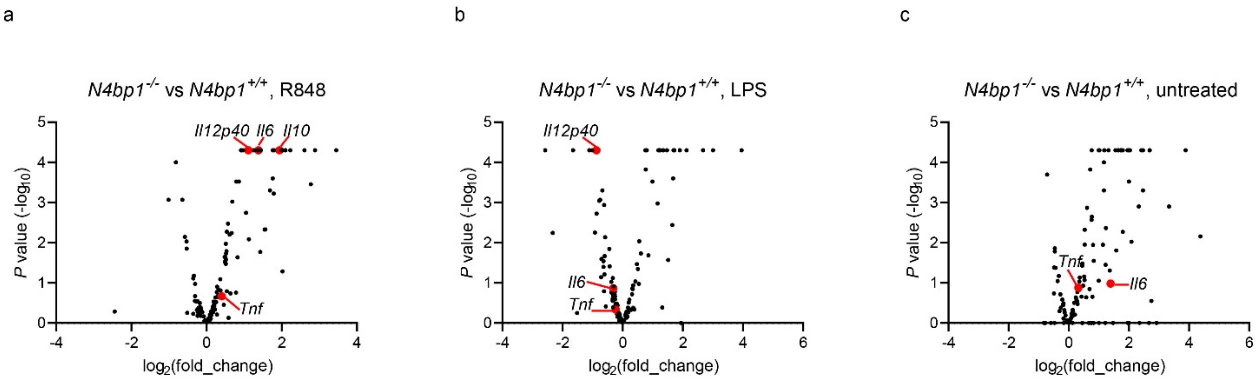


**Supplementary Figure 3. RNA_seq for *N4bp1*$^{+/+}$ and *N4bp1*$^{-/-}$ peritoneal macrophages. a-c,** Comparative analysis of expression levels of 124 NF-κB-dependent genes measured by RNA sequencing in *N4bp1*$^{+/+}$ vs. *N4bp1*$^{-/-}$ peritoneal macrophages stimulated with R848 (**a**), LPS (**b**) or without any stimulation (**c**). Cells were collected from two mice for each condition. *Il6* and *Tnf* were highlighted in red.

Supplementary Figure 4

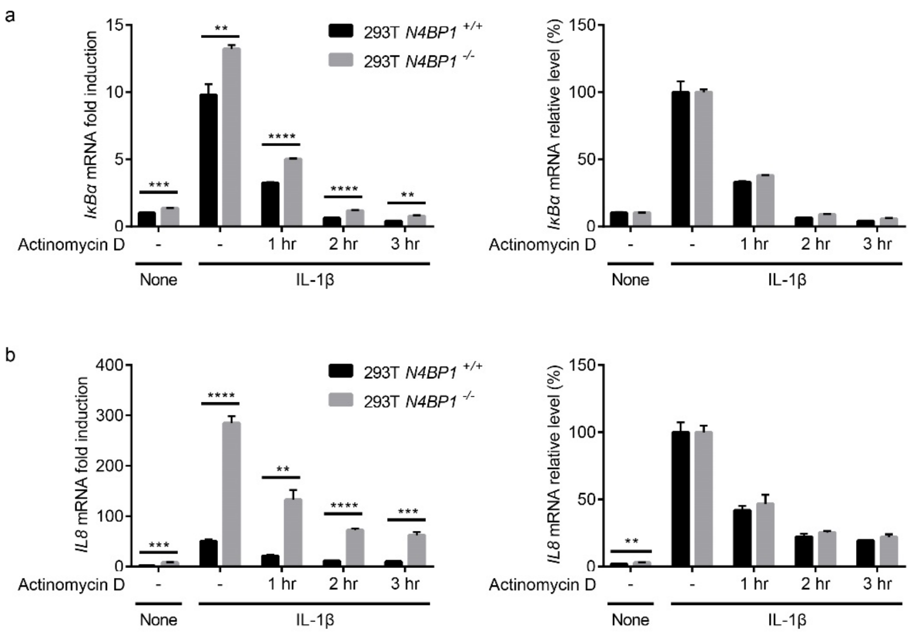


**Supplementary Figure 4. Comparable mRNA degradation rates in wild-type and *N4BP1$^{-/-}$* HEK293T cells. a, b,** RT-qPCR analysis of *NFKBIA* (*IκBα*) (**a**) or *IL8* (**b**) in wild-type and *N4BP1$^{-/-}$* HEK293T stimulated with IL-1β for 2 h then treated without or with actinomycin D (2 μg/ml) for the indicated times. *Left*, mRNA fold induction. *Right*, amount of mRNA relative to that of cells stimulated with IL-1β but without Actinomycin D, set as 100%. * $P < 0.05$, ** $P < 0.01$, *** $P < 0.001$, **** $P < 0.0001$ (two-tailed Student's *t*-test). Data are representative of two independent experiments (mean ± s.d.).

Supplementary Figure 5

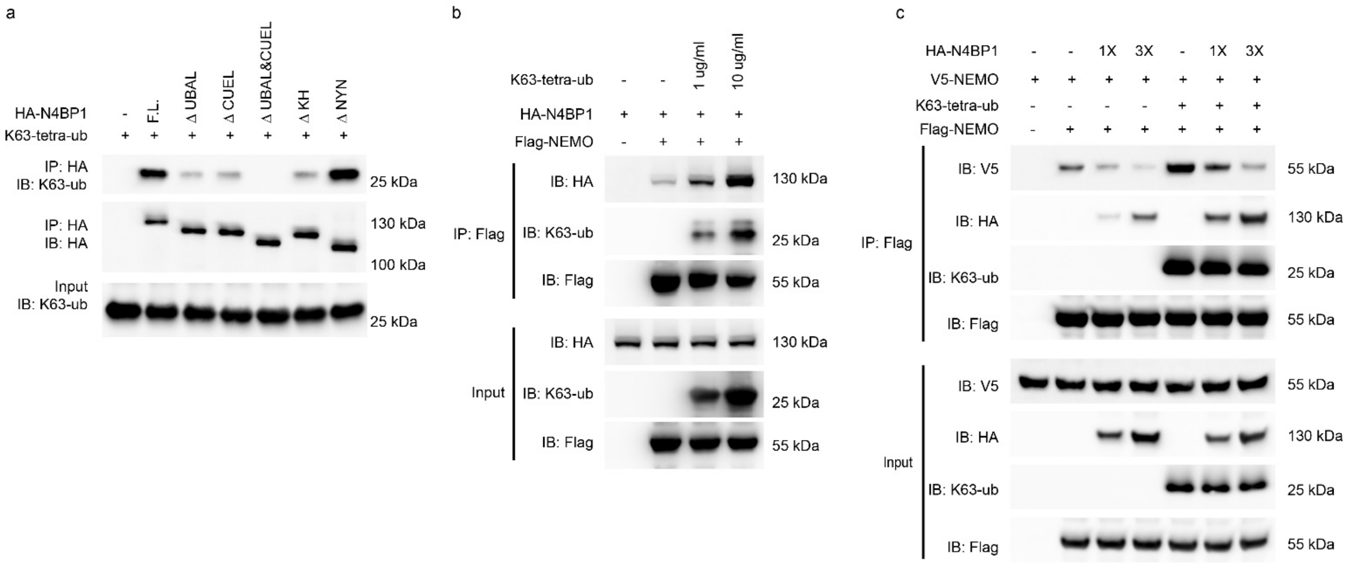


**Supplementary Figure 5. K63-linked ubiquitin enhances N4BP1 and NEMO interaction. a,** K63-linked ubiquitin was incubated without (-) or with purified recombinant HA-N4BP1 or its domain deletion forms and subjected to immunoprecipitation with anti-HA. **b,** Purified recombinant HA-N4BP1 and Flag-NEMO were incubated without (-) or with K63-linked ubiquitin as indicated, and subjected to immunoprecipitation with anti-Flag. **c,** Purified recombinant Flag-NEMO and V5-NEMO were mixed together and then incubated with different amounts of HA-N4BP1 in the presence of K63-linked ubiquitin or without ubiquitin, and subjected to immunoprecipitation with anti-Flag. Complexes were analyzed by immunoblotting with the indicated antibodies (**a-c**). Data are representative of two independent experiments.

Supplementary Figure 6

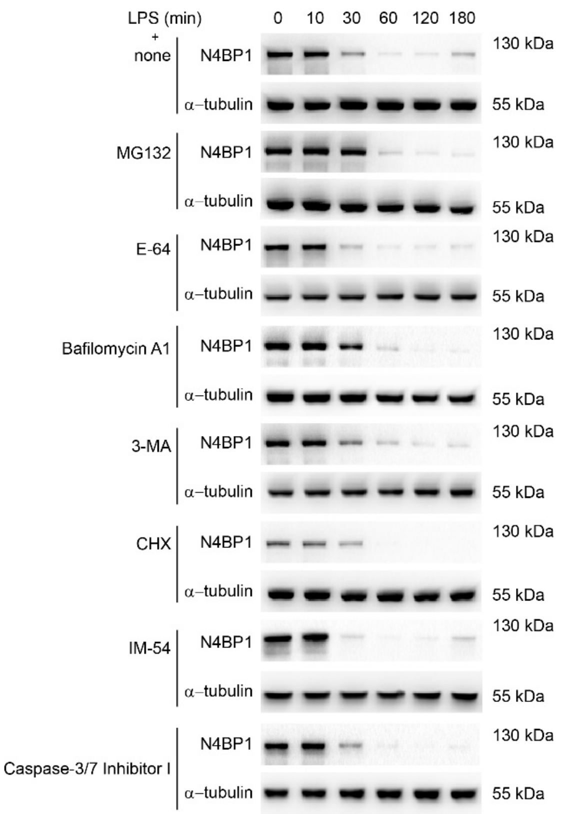


**Supplementary Figure 6. Downregulation of N4BP1 is not mediated by the proteasome, autophagy, or cysteine proteases.** Immunoblot analysis of N4BP1 in peritoneal macrophages pretreated with MG132 (10 μM), E-64 (20 μM), bafilomycin A1 (1 μM), 3-MA (5 mM), IM-54 (50 μM), or caspase-3/7 Inhibitor I (100 μM) for 1 h, or with CHX (50 μg/ml) for 3 h then stimulated with LPS for the indicated times. Data are representative of two independent experiments.

Supplementary Figure 7

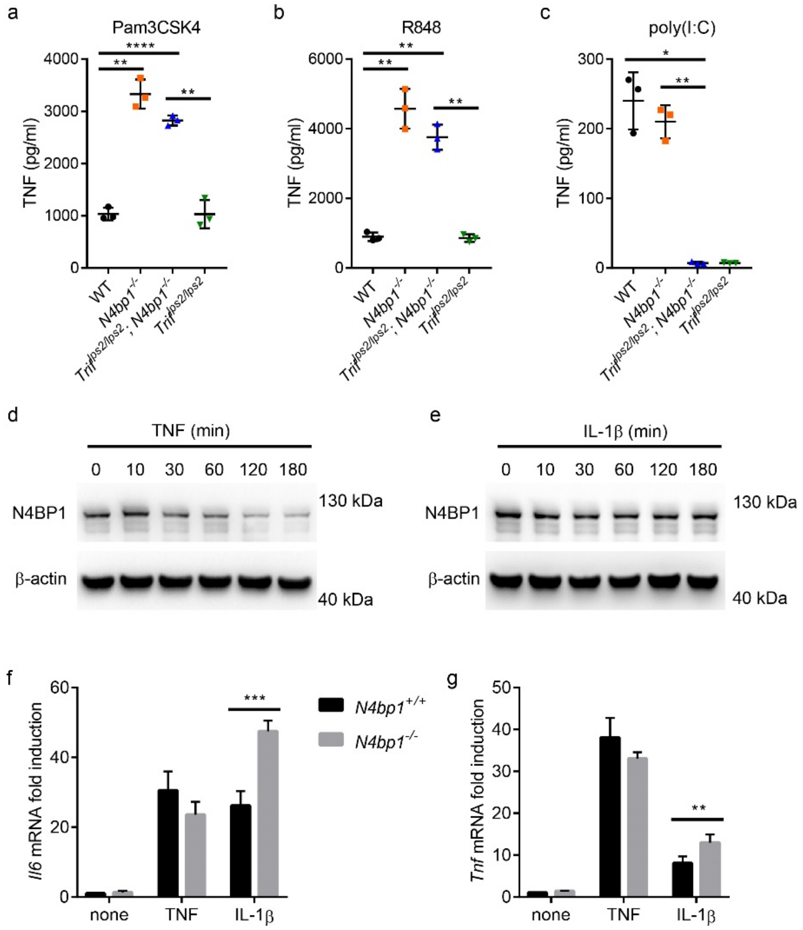


**Supplementary Figure 7. *N4bp1* deficiency enhanced IL-1β but not TNF signaling. a-c,** TNF concentration in the culture medium of wild-type (WT), $N4bp1^{-/-}$, $Trif^{lps2/lps2}N4bp1^{-/-}$, and $Trif^{lps2/lps2}$ peritoneal macrophages treated with Pam3CSK4 (**a**), R848 (**b**), or poly(I:C) (**c**). **d, e,** Immunoblot analysis of N4BP1 in peritoneal macrophages treated with TNF (100 ng/ml) (**d**) or IL-1β (100 ng/ml) (**e**). **f, g,** RT-qPCR analysis of *Il6* (**f**) or *Tnf* (**g**) in $N4bp1^{+/+}$ and $N4bp1^{-/-}$ peritoneal macrophages stimulated with TNF or IL-1β for 2 h. Each symbol (**a-c**) represents an individual mouse. * $P < 0.05$, **$P < 0.01$, ***$P < 0.001$, ****$P < 0.0001$ (two-tailed Student's *t*-test). Data are representative of two independent experiments (mean ± s.d. in **a-c,f,g**).
